# Repeated stimulation or tonic-signaling chimeric antigen receptors drive regulatory T cell exhaustion

**DOI:** 10.1101/2020.06.27.175158

**Authors:** Caroline Lamarche, German E. Novakovsky, Christopher N. Qi, Evan W. Weber, Crystal L. Mackall, Megan K Levings

**Affiliations:** Department of Surgery, University of British Columbia, Vancouver, BC, Canada; BC Children’s Hospital Research Institute, Vancouver, BC, Canada; Department of Medical Genetics, University of British Columbia, Vancouver, BC, Canada; Department of Pediatrics and Medicine, Stanford University School of Medicine, Stanford, CA, USA; Stanford Cancer Institute, Stanford University School of Medicine, Stanford, CA, USA; School of Biomedical Engineering, University of British Columbia, Vancouver, BC, Canada

## Abstract

Regulatory T cell (Treg) therapy is a promising approach to improve outcomes in transplantation and autoimmunity. In conventional T cell therapy, chronic stimulation can result in poor *in vivo* function, a phenomenon termed exhaustion. Whether or not Tregs are also susceptible to exhaustion, and if so, if this would limit their therapeutic effect, was unknown. We studied how two methods which induce conventional T cell exhaustion – repetitive stimulation or expression of a tonic-signaling chimeric antigen receptor (CAR) – affect human Tregs. With each repetitive polyclonal stimulation Tregs progressively acquired an exhausted phenotype, and became less suppressive *in vitro*. Tregs expressing a tonic-signaling CAR rapidly acquired an exhausted phenotype and had major changes in their transcriptome and metabolism. Although tonic-signaling CAR-Tregs remained stable and suppressive *in vitro*, they lost *in vivo* function, as tested in a model of xenogeneic graft-versus-host disease. The finding that human Tregs are susceptible to exhaustion has important implications for the design of Treg adoptive immunotherapy strategies.

## Introduction

Regulatory T cells (Tregs) control immune homeostasis and their adoptive transfer is a promising therapeutic approach with multiple trials completed, on-going or planned to test their efficiency in autoimmunity, solid organ transplantation or hematopoietic stem cell transplantation (reviewed in (Ferreira et al., 2019)). As Tregs are relatively rare, the majority of cell therapy protocols involve *in vitro* stimulation and expansion, in some cases culturing cells for up to 36 days to achieve > 2,000 fold expansion (MacDonald et al., 2019b). Although these *in vitro*-expanded cells typically remain FOXP3^+^ and suppressive *in vitro*, whether or not repeated stimulation of Tregs affects their *in vivo* function is unknown.

In contrast to the paucity of knowledge regarding how chronic stimulation affects Treg function, the effects of chronic *in vitro* or *in vivo* stimulation of T cells has been extensively studied. For example, in the context of infection, cancer or transplantation, chronic exposure to antigen leads to a phenomenon termed exhaustion, characterized by high expression of inhibitory receptors such as programmed cell death protein 1 (PD-1), T-cell immunoglobulin and mucin-domain containing-3 (TIM-3), and lymphocyte-activation gene 3 (LAG-3), diminished proliferative and cytokine-producing capacity, and high apoptosis (Fribourg et al., 2019; Saeidi et al., 2018; Thorp et al., 2015; Wherry and Kurachi, 2015). Although most research in this area examined effects on CD8^+^ T cells or unfractionated CD3^+^ T cells, but there is evidence that CD4^+^ T cells are similarly susceptible to exhaustion (Lynn et al., 2019; Shahbaz et al., 2020; Zou et al., 2020).

Exhaustion can also be induced by expression of engineered receptors, including chimeric antigen receptors (CARs), which mediate tonic signaling via spontaneous aggregation in the absence of antigen (Long et al., 2015; Lynn et al., 2019; Weber et al., 2020b). CAR-mediated T cell exhaustion induces expression of a variety of inhibitory receptors and diminishes their anti-tumor therapeutic efficacy (Finney et al., 2019; Wherry and Kurachi, 2015). This state of functional exhaustion can be manipulated by limiting activity of signaling pathways, and altering the epigenetic program and/or metabolic pathways (Chen et al., 2019; Fraietta et al., 2018; Lynn et al., 2019; Weber et al., 2020b).

Whether or not Tregs are also susceptible to exhaustion has not been explored. Here we used two different methods to chronically stimulate human Tregs and investigated the phenotypic, metabolic and functional consequences.

## Results

### Repetitive polyclonal Treg stimulation leads to decreased suppressive function

Tregs are relatively rare cells that are hypo proliferative, so most protocols to expand Tregs for adoptive immunotherapy typically involve at least two polyclonal stimulations with anti-CD3 and -CD28 mAbs to obtain clinically relevant numbers (MacDonald et al., 2019b). To ask if repetitive stimulation could be detrimental for Treg function, we polyclonally stimulated Tregs weekly for 4 weeks and analyzed changes in their phenotype and function over time. Experiments were done with naïve CD4^+^CD25^hi^CD27^low^CD45RA^+^ Tregs (**Supp Figure 1A**) to minimize contamination with CD4^+^ conventional T cells (Tconvs) and/or unstable Tregs (Hoffmann et al., 2006). We first examined expression of cell surface receptors associated with exhausted T cells (Wherry and Kurachi, 2015), and found repeated Treg stimulation resulted in progressively higher expression of LAG-3 and TIM-3, but not of PD-1 (**Figure 1A)** or glucocorticoid-induced TNFR-related protein (GITR**) (Figure 1B)**.

**Figure 1.**
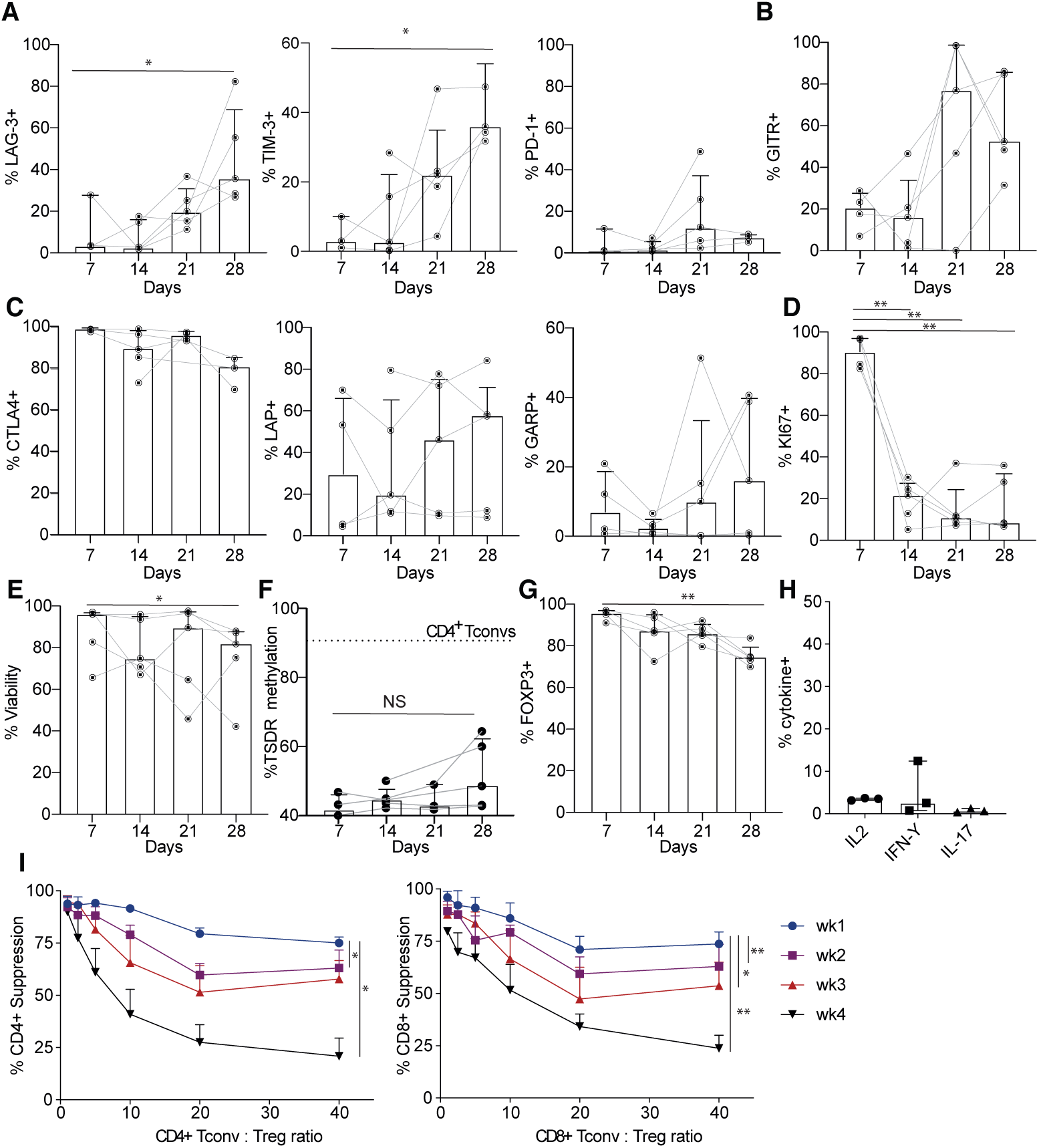
Repetitive stimulation of Tregs leads to dysfunction. Naïve Tregs were sorted and stimulated weekly for 4 weeks. Analysis for (**A**) LAG-3, TIM-3, PD-1, **(B)** GITR, (**C**) CTLA4, LAP, and GARP, (**D**) Ki67 and (**E**) viability were assessed weekly. (**F**) DNA was isolated and pyrosequencing was used to measure methylation status of 7 CpGs in the Treg specific demethylation region (TSDR), only females were used for these experiments. For reference, the average amount of methylation in CD4^+^ conventional T cells (Tconv) is indicated by the dotted line. **(G)** Expression of FOXP3 was assessed weekly. (**H**) Intracellular cytokine staining was done after 28 days of culture. n=3 from 2 independent experiments (**I**) Cell-proliferation dye (CPD)450-labelled PBMCs were stimulated with anti-CD3/28 Dynabeads in the absence or presence of the indicated ratio of Tregs at the end of each weekly re-stimulation. After 4 days, the division index of gated CD4^+^ or CD8^+^ T cells was determined and percent suppression was calculated. To compare suppression over time, suppression assays used the same responder PBMC using replicate cryopreserved aliquots. **(A-H)** Mixed-effects model analyses were done with Dunnett’s multiple comparisons test to compare changes overtime to day 7. Each dot represents a unique donor. Median ± interquartile range, n=3-5 from 3 independent experiments. (**I**) One-way ANOVA with Dunnett’s multiple comparisons test, mean ± SEM, n=4-5 from 2-3 independent experiments. ** p ≤ 0.01, * p ≤ 0.05.

We then measured cell-surface proteins associated with Treg function, finding that the naturally high expression of cytotoxic T-lymphocyte-associated protein 4 (CTLA-4) was not further enhanced by repeated stimulation. There was also no effect on expression of latency-associated peptide (LAP, inactive TGF-*β*), or glycoprotein A repetitions predominant (GARP, a receptor for LAP) (**Figure 1C**). However, as seen with exhausted T cells (Wherry, 2011), repeatedly-stimulated Tregs became progressively less proliferative (decrease in % Ki67^+^ cells) and viable (**Figure 1D&E**).

To ask if these changes could be related to changes in Treg stability, we measured methylation at the Treg-specific demethylation region of *FOXP3*, finding no significant increase over time (**Figure 1F)**. Similarly, FOXP3 expression was stable until the final re-stimulation when the average proportion of FOXP3^+^ cells declined to 75±5% at day 28 (**Figure 1G**).

Consistent with preserved Treg stability, production of inflammatory cytokines remained low (**Figure 1H)**. Nevertheless, as early as the second stimulation, the Tregs became progressively less suppressive as judged by the significant decrease in their ability to suppress division of CD4^+^ or CD8^+^ T cells (**Figure 1I**). Overall, these data indicate that repetitive, chronic stimulation of Tregs causes increased expression of some proteins characteristic of exhausted T cells and is deleterious for their *in vitro* function.

### Tonic signaling CARs induce a phenotype consistent with exhaustion in Tregs

To further study the existence and biological relevance of Treg exhaustion, we next investigated the effect of expressing a tonic signaling CAR (TS-CAR), which incorporates a high-affinity scFv specific for disialoganglioside (GD2) fused to a CD28 co-stimulatory endodomain and CD3*ζ* (Long et al., 2015; Lynn et al., 2019; Weber et al., 2020a). Expression of this TS-CAR induces hallmark functional, phenotypic, transcriptional, and epigenetic features of exhaustion in T cells *in vitro* and abrogates antitumor effects *in vivo* (Long et al., 2015; Lynn et al., 2019). Naïve CD4^+^CD127^lo^CD25^hi^ CD45RA^+^CD62L^hi^ Tregs, or as a control naïve CD4^+^ CD45RA^+^CD62L^hi^ Tconvs (**Supp Figure 1A**), were untransduced or transduced with the TS-CAR, or a CD19-specific control CAR (non-TS-CAR) which neither mediates tonic signaling nor induces exhaustion (Long et al., 2015; Lynn et al., 2019). After 7 days, CAR-expressing cells were sorted then cultured for an additional 5 days before analysis. Phenotypic analysis revealed that, in comparison to untransduced or non-TS-CAR Tregs, a significantly higher proportion of TS-CAR Tregs expressed inhibitory receptors associated with exhaustion, namely LAG-3, TIM-3, PD-1 **(Figure 2A and Supp Figure 1B**) as well as GITR **(Figure 2B and Supp Figure 1B)**. Similarly, a higher proportion of TS-CAR Tregs expressed the Treg-function-associated proteins CTLA-4, LAP and GARP **(Figure 2C and Supp Figure 1B)**. High expression of all these proteins was also observed in TS-CAR-expressing CD4^+^ Tconvs and CD8^+^ T cells (**Supp Figure 2A**). Contrary to our observations with repeatedly stimulated Tregs, a higher proportion of TS-CAR Tregs and CD4^+^ Tconvs were Ki67^+^ **(Figure 2D and Supp Figure 2B**), which could be indicative of a high proportion of terminally exhausted cells as opposed to progenitor exhausted cells (Miller et al., 2019). Although viability was similar between the 3 groups, in accordance with exhausted T cells (Zhang et al., 2020), TS-CAR Tregs were more apoptotic than controls **(Figure 2E)**.

**Figure 2.**
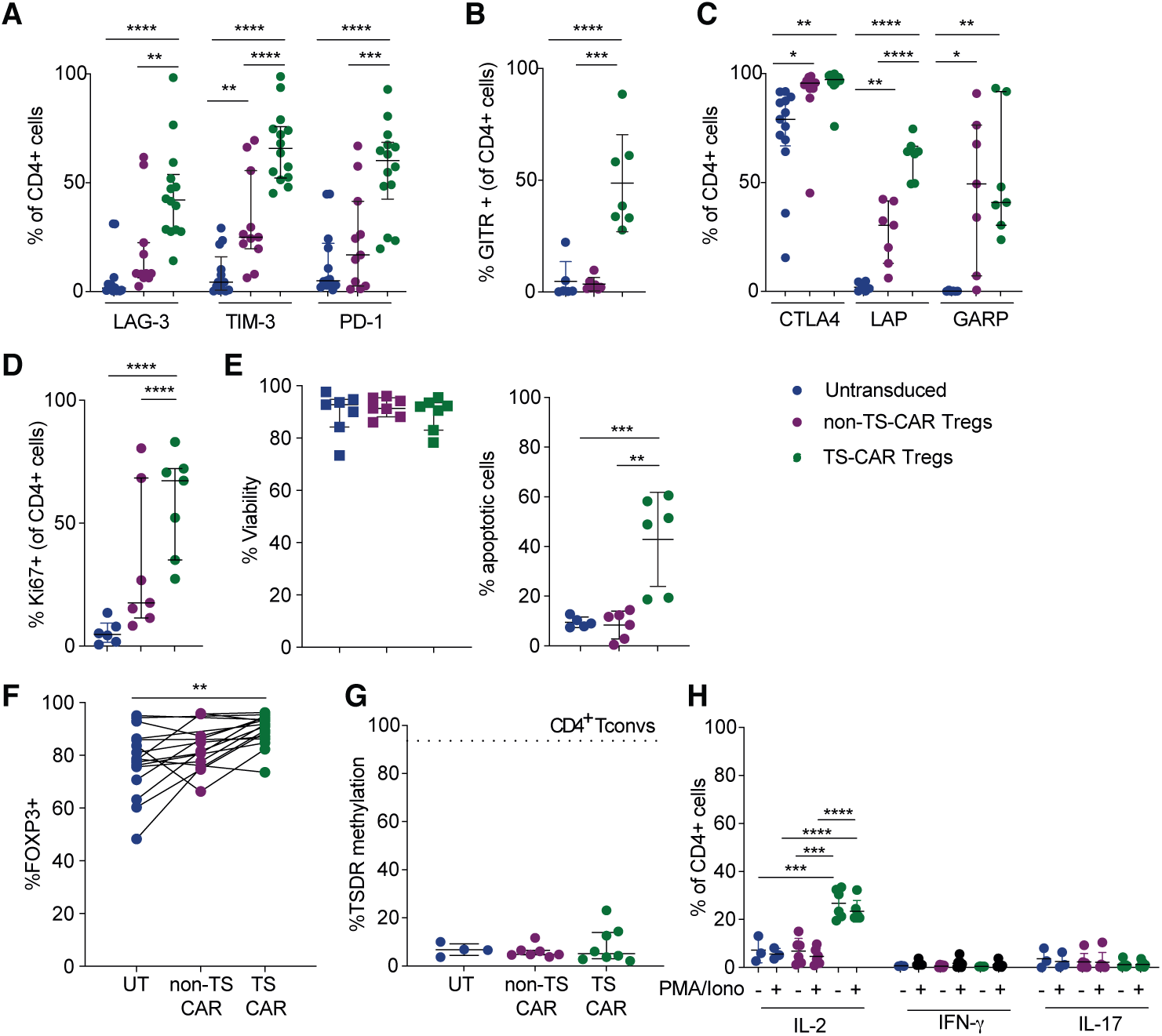
Expression of a tonic signaling CAR in Tregs induces phenotypic changes without lineage destabilization. Naïve Tregs were left transduced (UT) or transduced with retrovirus encoding a non-tonic signaling (non-TS) or a tonic signaling (TS-) CAR. After 11-12 days of culture cells were analyzed for expression of: **(A)** LAG-3, TIM-3, PD-1, **(B)** GITR, **(C)** CTLA4, LAP, GARP, and **(D)** Ki67. **(E)** Viability was assess using an automated cell counter and apoptosis by flow cytometry and **(F)** FOXP3 by intracellular staining. **(G)** DNA was isolated and pyrosequencing was used to measure methylation status of 7 CpGs in the Treg specific demethylation region (TSDR). Data are the average methylation of the 7 CpGs. Only males were used for these experiments. For reference, the average amount of methylation in CD4^+^ conventional T cells (Tconv) is indicated by the dotted line. (**H)** Intracellular cytokine production was assessed by flow cytometry 4 hours after a PMA/Ionomycin stimulation. Each dot represents a unique donor. Median ± interquartile range. (**A-H**) One-way Anova with Turkey’s comparisons test. For **A-E** (left), n=11-14 from 5-6 independent experiments. For the apoptosis assay (**E)**, right, n=5-6 from 3-4 independent experiments. For **F**, n=11-17 from 5-7 independent experiments. For **G**, n=3-6 from 3-4 independent experiments. For **H**, n=3-6 from 3 independent experiments. **** p ≤ 0.0001, *** p ≤ 0.001, ** p ≤ 0.01, * p ≤ 0.05.

These phenotypic changes were not associated with loss of Treg stability, since TS-CAR Tregs had no change in TSDR methylation and actually increased FOXP3 expression (**Figure 2 F&G**). Nevertheless, compared to controls, a significantly higher proportion of TS-CAR Tregs spontaneously produced IL-2 (**Figure 2H**), suggesting that these cells overcame FOXP3-mediated repression of *IL2* transcription (Bettelli et al., 2005). Overall, these data show that Tregs expressing a TS-CAR acquire a phenotype which mirrors that acquired by exhausted CD8^+^ T cells and CD4^+^ Tconvs (**Supp Figure 2**) (Lynn et al., 2019).

### Transcriptome analysis of tonic signaling CAR-Tregs reveals changes in metabolic pathways

A limitation of studying exhaustion in Tregs is that several of the canonical exhaustion makers are also markers of Tregs, or simply of activated T cells (Tanaka and Sakaguchi, 2019; Wherry and Kurachi, 2015). Moreover, since Tregs do not produce inflammatory cytokines, loss of cytokine production cannot be monitored as a surrogate marker of dysfunction. Therefore, we next took an unbiased approach and used RNA sequencing to determine how tonic signaling affected the Treg transcriptome in comparison to untransduced or non-TS-CAR Tregs, or CD4^+^ Tconvs. Multi-dimensional scaling plots revealed that Tregs and Tconvs clustered separately, and that, for both Tregs and Tconvs, the TS-CARs were further away from untransduced cells than cells expressing the non-TS-CAR cells (**Figure 3A**).

**Figure 3.**
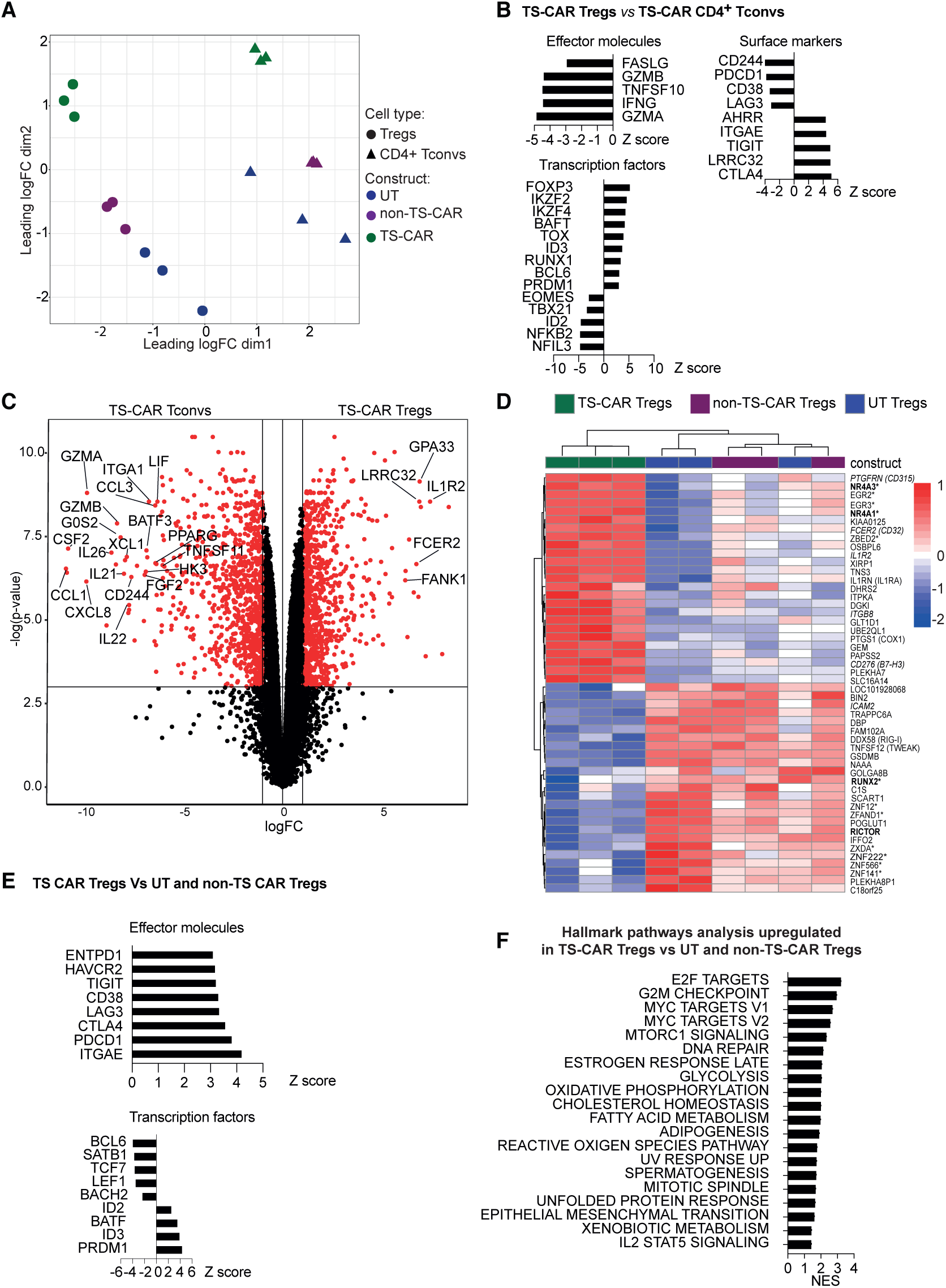
Expression of a tonic signaling CAR results in differential effects on the transcriptome of Tregs and Tconvs. After 12 days of culture, RNA was extracted from Tregs or CD4^+^ Tconvs that were untransduced (UT) or transduced to express a non-tonic signaling (non-TS) or a tonic signaling (TS-) CAR, and subject to RNA sequencing. **(A)** Multidimensional scaling plots comparing the transcriptome of the indicated types of Tregs and Tconvs. **(B)** Z score of differentially expressed genes in TS-CAR Tregs compared to UT or non-TS-CAR Tregs. Threshold used was a logFC > 1 and adj.pvalue < 0.05, and logFC < −1 and adj.pvalue < 0.05. **(C)** Volcano plot comparing the gene expression profile of TS-CAR Tregs versus TS-CAR Tconvs after removal of genes differentially expressed (upregulated and downregulated) between UT Tconvs and UT Tregs (see Supp. Figure 3A). Genes in red are defined by a p-value < 0.05. **(D)** Heatmap showing the top 50 differentially expressed genes between the different types of Tregs. Bold genes are associated with exhaustion in conventional T cells, italicized genes are cell surface proteins and those marked with * are transcription factors. **(E)** Z score comparing genes that were significatively upregulated or downregulated in TS-CAR Tregs versus TS-CAR Tconvs. Threshold used was a logFC > 1 and adj.pvalue < 0.05, and logFC < −1 and adj.pvalue < 0.05. **(F)** Gene set enrichment analysis. Normalized enrichment score (NES) of hallmark pathways analysis overrepresented in TS-CAR Tregs compared to non-TS CAR or UT Tregs cells. n=3 per group from 2 independent experiments.

To better understand the consequences of exhaustion in Tregs, we first sought to identify differences in gene expression between TS-CAR Tregs and TS-CAR CD4^+^ Tconvs. As expected, in comparison to TS-CAR CD4^+^ Tconvs, TS-CAR Tregs had low expression of effector molecules such as *GZMA, GZMB* (granzyme A &B), *IFNG, TNFSF10* (TRAIL) and *FASLG*, and high expression of Treg-related genes such as *FOXP3, IKZF2* (Helios) and *LRRC32* (GARP), confirming their Treg identity (**Figure 3B**). Interestingly, inhibitory surface receptors were differentially expressed in TS-CAR Tregs compared to TS-CAR CD4^+^ Tconvs, with lower expression of *CD244, PDCD1* (PD1) and *LAG3* and higher expression of *TIGIT* and *CTLA4*. Similarly, there were differences in expression of exhaustion-related transcription factors (Zhang et al., 2020), with higher expression of *TOX*, but not *EOMES* or *TBX21* in TS-CAR Tregs compared to TS-CAR CD4^+^ T cells (**Figure 3B**).

Since many of the genes differentially expressed between TS-CAR Tregs and Tconvs were simply characteristic of known transcriptional differences between these cell lineages, we also examined differential gene expression after removing genes that were differentially expressed between UT Tregs and UT Tconvs (**Figure 3C** and **Supp Figure 3A**). The resulting data revealed that compared to CD4^+^ TS-CAR Tconvs, TS-CAR Tregs expressed significantly less *BATF3*, a transcription factor expressed in exhausted TS-CAR T cells (Lynn et al., 2019; Weber et al., 2020b) and less *CD244*, an immunoregulatory receptor also associated with T cell exhaustion (Agresta et al., 2018; Schlaphoff et al., 2011). TS-CAR Tconvs also expressed more chemokines (*CCL1, CXCL1* and *CXCL8*), cytokines (*IL21, IL22* and *LIF*) and integrin *α*1 (*ITGA1*). On the other hand, TS-CAR Tregs overexpressed *GPA33*, the expression of which is a marker of stable Tregs (Opstelten et al., 2020), and GARP (*LLRC32*), confirming the notion that tonic signaling in Tregs does not impair their lineage stability.

Within the 3 groups of Tregs, the transcriptome of the TS-CAR Tregs was clearly distinct from UT and non-TS-CAR Tregs, with a heat map of the 50 most differentially expressed genes between these groups shown in **Figure 3D**. In comparison to both UT and non-TS CAR Tregs, TS-CAR Tregs had high expression of *NR4A3* (NOR1) and *NR4A1 (*NUR77), two genes associated with exhaustion in CD8^+^ T cells (Chen et al., 2019; Liu et al., 2019). TS-CAR Tregs also had lower *RICTOR*, which is part of the mammalian target of rapamycin (mTOR) complex 2 (mTORC2) and implicated in control of Treg suppressive function (Charbonnier et al., 2019). There was also a decrease in *RUNX2*, the expression of which is enhanced by a weak rather than a strong TCR stimulation (Olesin et al., 2018). Confirming the flow cytometry data, TS-CAR Tregs had high expression of exhaustion-related genes such as *PDCD1* (PD1), *HAVCR2* (TIM3), *TIGIT, LAG3* and *CTLA4*, but low expression of several transcription factors known to be downregulated in terminally exhausted T cells, including *BCL6, SATB1, TCF7* (which encodes TCF-1), *LEF1* and *BACH2* (Blank et al., 2019; Miller et al., 2019; Wang et al., 2019) **(Figure 3E)**.

Hallmark pathway analysis, obtained by Gene Set Enrichment Analysis, revealed that TS-CAR Tregs upregulated pathways linked to metabolism, including MYC and MTORC1 signaling, glycolysis, oxidative phosphorylation, cholesterol homeostasis, fatty acid metabolism and adipogenesis (**Figure 3F**), suggesting that changes in cell metabolism may be key factors driving the observed phenotypic changes in TS-CAR Tregs. However, although these pathways were upregulated in TS-CAR Tregs compared to other Treg groups, they were still lower in comparison to TS-CAR CD4^+^ Tconvs (**Supp Figure 3B**).

### Tonic signaling in Tregs causes an increase in glycolysis

A growing body of evidence suggests that differences in metabolic pathway activity underlie functional differences between Tregs and Tconvs (Beier et al., 2015). Specifically, oxidative phosphorylation and mitochondrial activity, which are required for Treg suppressive function (Pacella and Piconese, 2019; Saravia et al., 2020), are promoted by FOXP3 via inhibition of c-Myc-driven glycolysis (Angelin et al., 2017; Gerriets et al., 2016; Howie et al., 2017; Wang et al., 2011). In the context of exhaustion, metabolic dysfunction is known to precede its development in T cells (McKinney and Smith, 2018), and is influenced by CAR expression (Kawalekar et al., 2016). Although FOXP3 expression was similar between UT, non-TS-and TS-CAR Tregs, transcriptomic analysis suggested that there could be TS-CAR related changes in metabolic activity. Therefore, we analyzed their extracellular acidification rate (ECAR) and oxygen consumption rate (OCR) and as surrogate measures of glycolysis and oxidative phosphorylation, respectively. In their basal state TS-CAR Tregs had a tendency to have higher ECAR and OCR than UT or non-TS-CAR Tregs, suggesting both metabolic pathways were enhanced (**Figure 4A&B, left**). However, a significantly lower relative OCR/ECAR ratio showed that TS-CAR Tregs preferentially engaged glycolysis compared to their UT or non-TS-CAR counterparts (**Figure 4C**).

**Figure 4.**
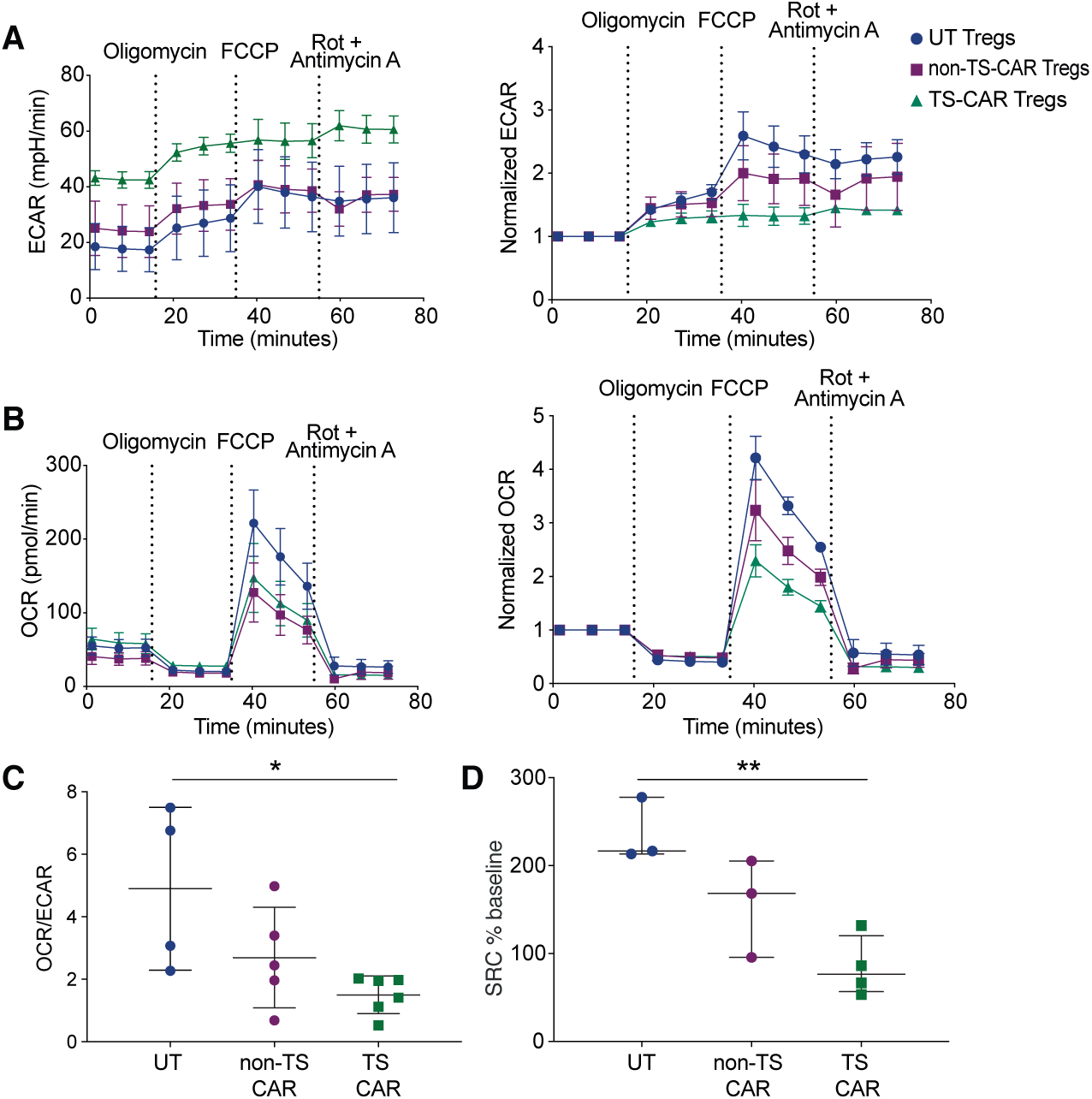
Expression of a tonic signaling CAR signaling alters Treg metabolism. Tregs were untransduced (UT) or transduced to express a non-tonic signaling (non-TS) or a tonic signaling (TS-) CAR. After 11-12 days of culture cells were plated in Cell-Tak coated wells and subject to the Agilent Seahorse XF Cell Mito Stress Test. **(A)** Extracellular acidification rate (ECAR) (left) and normalized to baseline ECAR (right) over time, **(B)** Oxygen consumption rate (OCR) (left) and normalized to baseline OCR (right) over time. Oligomycin, FCCP and a mix of Rotenone and antimycin A were added as indicated. **A&B)** n=3-5 donor from 2 independent experiments, using the average of technical replicates. Mean ± SEM. **(C)** OCR/ECAR ratio at baseline. One-way Anova with Turkey’s comparisons test p=0.0196. n=4-6 from 3 independent experiments. (**D**) The spare respiratory capacity (SRC) was calculated by the average of the maximum OCR after FCCP injection minus the average basal respiration, divided by the average basal respiration, times 100 ((max-basal)/basal*100). One-way Anova with Turkey’s comparisons test p=0.0054, n=3-4 from 2 independent experiments.

To further interrogate these pathways, cells were exposed to oligomycin, a mitochondrial ATP synthase inhibitor which prevents oxidative phosphorylation and usually upregulates glycolysis. Since TS-CAR Tregs had high basal ECAR, the relative oligomycin-stimulated increase was modest compared to the other groups, suggesting limited spare glycolytic capacity (**Figure 4A**). In contrast there was comparable ATP-coupled mitochondrial respiration in all groups (**Figure 4B**). Injection of carbonyl cyanide-4 (trigluoromethoxy) phenylhydrazone (FCCP) to maximize mitochondrial oxygen consumption, revealed that TS-CAR Tregs also had lower spare respiratory capacity (**Figure 4D**). Data in Tregs were mirrored with CD4^+^ Tconvs (**Supp Figure 3C**). Overall these data suggest that TS-CAR Tregs would poorly adapt to an increase in energy demand due to their high basal activity of both of these pathways (Schurich et al., 2016; van der Windt et al., 2012).

### TS-CAR Tregs are dysfunctional *in vivo* but not *in vitro*

In mice upregulation of glycolysis and/or inhibition of oxidative phosphorylation leads to reduced Foxp3 expression and/or suppressive function (Beier et al., 2015; Gerriets et al., 2016; Wei et al., 2016). We therefore next asked how TS-CAR expression affected Treg function. We first tested suppressive function *in vitro* by co-culturing increasing numbers of UT, non-TS-, or TS-CAR Tregs with anti-CD3/28-stimulated PBMCs and measuring proliferation after 4 days. Unexpectedly, TS-CAR Tregs were significantly more potent than UT or non-TS Tregs at suppressing CD4^+^ and CD8^+^ T cell proliferation (**Figure 5A**), but within the Treg population, the TS-CAR Tregs proliferated significantly less than UT or non-TS-CAR Tregs (**Figure 5B**). The increase in TS-CAR Tregs suppressive function despite their lower division index could be due to a higher suppression potential on a per cell basis and/or by the immunomodulatory potential of apoptotic Tregs (Maj et al., 2017).

**Figure 5.**
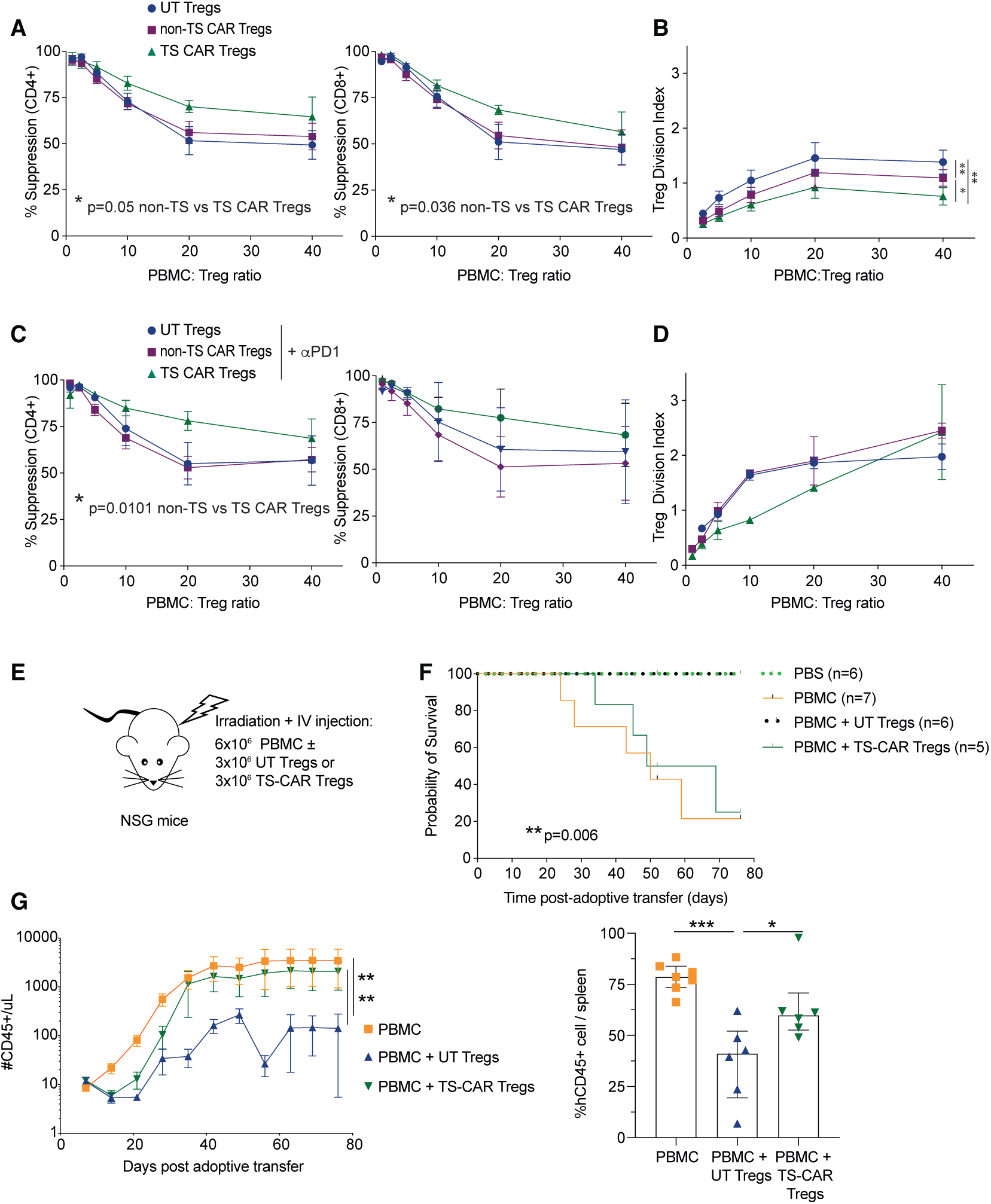
Tonic signaling CAR Tregs are functional in vitro but not in vivo. **(A)** Suppression of CPD-ef450-labeled PBMCs by untransduced (UT), non-tonic signaling (non-TS) or tonic signaling (TS-) CAR CPD-ef-670-labeled Tregs was determined after 4 days of co-culture. n=8-10 from 4 independent experiments **(B)** The Treg division index from cultures in (A) was determined. n=6-7 from 3 independent experiments **(C &D)** Suppression assays were done with the addition of nivolumab (anti-PD-1 mAb) at 10mg/ml. **(A-D)** Analysis were done using one-way ANOVA with Turkey’s multiple comparisons test. **(E-G)** Irradiated NSG mice were injected with PBS or 6×10^6^ PBMCs without or with 3×10^6^ UT or TS-CAR Tregs. Data from 2 independent experiments (**E)** Schematic diagram of the experiment set up. **(F)** Survival curve, log-rank (Mantel-Cox) test. **(G)** Absolute number of human CD45+ cells/μL of blood assessed weekly after adoptive transfer. One-way ANOVA with Dunnett’s multiple comparisons test (left) and % Human CD45+ engraftment in the spleen upon experimental or humane endpoint (of live singlets) in the different groups (right). One-way ANOVA with Turkey’s multiple comparisons test.

To address why expression of a TS-CAR increased, whereas repetitive polyclonal stimulation decreased (**Figure 1H**) in vitro suppression, we noted the difference in PD-1 expression. Specifically, PD-1 was highly expressed by TS-CAR Tregs but not by repetitively stimulated Tregs. To investigate if the increased in vitro suppressive activity of TS-CAR Tregs was mediated by PD-1, suppression assays were repeated in the presence of nivolumab, an inhibitory anti-PD-1 mAb. Although nivolumab did not reverse the heightened suppressive activity of TS-CAR Tregs (**Figure 5C**), it reversed the decreased TS-CAR Treg-intrinsic proliferation (**Figure 5D**). The beneficial effect of PD-1 inhibition on Treg proliferation is consistent with the finding that PD-1 blockade in patients can sometimes lead to paradoxical cancer progression due to enhanced Treg activity (Kamada et al., 2019).

Studies of TS-CARs in Tconvs showed that despite normal *in vitro* effector activity, functional defects were revealed *in vivo* (Long et al., 2015) so we next tested Treg function *in vivo* using a humanized mouse model of xenogeneic graft-versus-host disease. NSG mice were injected with 6×10^6^ PBMCs without or with 3×10^6^ UT or TS-CAR (*n*.*b*. non-TS Tregs not examined as they would be stimulated by CD19^+^ B cells in the PBMCs), monitored and bled weekly (**Figure 5E**). Surprisingly, TS-CAR Tregs completely lost their *in vivo* protective effect, with no improved survival in comparison to mice which only received PBMC. In contrast, mice which received UT Tregs were protected from mortality over this time period (**Figure 5F**).

Human cell engraftment mirrored the clinical effect, with only the UT Tregs able to reduce the number of circulating and splenic human CD45^+^ cells in comparison to mice that only received PBMCs (**Figure 5G**).

Continuous TCR signaling is essential for maintenance of immune tolerance by Tregs (Levine et al., 2014; Vahl et al., 2014), but whether or not Tregs become susceptible to exhaustion with prolonged, high level levels of stimulation was an outstanding question. We used two clinically-relevant approaches to ask whether exhaustion has the potential to limit the therapeutic efficacy of Tregs. With both repetitive polyclonal stimulation and TS-CAR expression we found clear evidence for development of a functional deficit consistent with the concept of exhaustion. Whereas repeated polyclonal stimulation gradually decreased *in vitro* suppression over time, expression of a TS-CAR caused rapid phenotypic and functional changes, and complete loss of suppression function *in vivo*. Importantly, loss of suppression function was not correlated with loss of Treg lineage stability, indicating that undesired acquisition of effector function is not a sequalae of chronic stimulation. These findings have important implications for the design of Treg adoptive transfer therapies and highlight the need to assess suppressive function both *in vitro* and *in vivo*.

A challenge to studying Treg exhaustion *in vitro* is the fact that inhibitory receptors characteristic of exhausted Tconvs are often associated with enhanced Treg suppressive potential (Anderson et al., 2016; Gautron et al., 2014; Gianchecchi and Fierabracci, 2018). For example, CTLA4 is essential for Treg function (Wing et al., 2008) and TIM-3^+^ Tregs are more potent than their TIM-3^-^ counterparts (Gautron et al., 2014). Moreover, as Tregs do not normally secrete inflammatory cytokines or exhibit cytolytic activity, *in vitro* analysis of these activities cannot be used to gauge this phenomenon. Indeed, repeated stimulation and TS-CAR expression increased expression of LAG-3 and TIM-3 but had little effect on other classical surface markers of exhaustion. Since cytometry-based methods typically used to evaluate exhaustion in T cells seem to have limited relevance in Tregs, it will of interest to study if cell surface proteins upregulated in TS-CAR Tregs at the mRNA level, including *FCER2* (CD32), *IL1R2* (IL1 decoy receptor), *CD276* (B7-H3) and *ITGB8* (integrin beta 8), could be used to monitor this state.

Gene expression analysis revealed high expression of exhaustion-related transcription factors, including *NR4A1* and *3, BATF* and *PRDM1* (Blimp1), suggesting there is at least some overlap with the transcriptional program that induces exhaustion in Tregs and in T cells. Moreover, as with exhausted T cells (Kawalekar et al., 2016; Wherry, 2011), exhausted Tregs displayed increased apoptosis, and underwent metabolic remodeling that reduced metabolic flexibility (Schurich et al., 2016). Poor metabolic flexibility in exhausted Tregs could underlie the difference in function *in vitro* versus *in vivo*: in comparison to *in vitro* cultures with saturated metabolite availability, limited availability of metabolic substrates *in vivo* could contribute to the profound loss of TS-CAR Treg function *in vivo*.

In mice, Tregs with a liver kinase B1 (Lkb1) deletion expressed a similar phenotype to TS-CAR Tregs with high expression of Pd1 and Gitr, increased apoptosis, and altered metabolism (Yang et al., 2017). Notably, Lkb1-deficient Tregs failed to suppress Th2 responses *in vivo*, an effect reversed by PD1 blockade (Yang et al., 2017). Although we did not observe decreased expression of *LKB1*, or of other genes found to mediate the “exhausted” phenotype in mouse Lkb1-deficient Tregs, including *CTNNB1* β-catenin and *HDC* (histidine decarboxylase), this pathway could still be affected due to alterations in downstream kinase mediators, and/or post transcriptional effects on LKB kinase activity. Investigation into how Treg exhaustion in humans might relate to this pathway, specifically with respect to effects on defined Th cell subsets, will be an important area of future investigation.

We noted similarities and differences between Tregs subject to chronic re-stimulation versus activity of a TS-CAR. In both cases there was increased expression of LAG-3 and TIM-3 and functional manifestations, but differential effects on proliferation, FOXP3 expression and *in vitro* suppressive function. The latter maybe driven by high expression of inhibitory receptors other than PD-1 by the TS-CAR Tregs. A better understanding of how such receptors contribute to Treg function in the context of exhaustion is important for understanding how immune checkpoint inhibitors might affect immune suppression in cancer and autoimmunity (Saleh and Elkord, 2019).

Finding ways to mitigate Treg exhaustion is critical for the continued development of successful Treg adoptive therapy. Our data highlight that repetitive stimulation should be limited during cell manufacturing, and that for optimal function, engineered receptors should be designed to limit antigen-independent activity. Our data also highlight to need to assess suppressive function *in vivo*, as similar to exhausted T cells, the design of typical *in vitro* assays may not fully reveal functional defects. An open question is how repeated exposure of a Treg to its target *in vivo* would affect long-term function. For example, even in the absence of tonic signaling, how continuous presence of an auto or alloantigen would affect a CAR-or TCR-engineered Tregs will be an important consideration. The metabolic phenotype of exhausted Tregs suggests that, as for CD8^+^ T cells and CD4^+^ T convs, strategies to modulate signaling pathway activity for example by use of kinase inhibitors (Dufva et al., 2020; Mestermann et al., 2019; Weber et al., 2019; Weber et al., 2020b), or modulation of transcription factor expression (Chen et al., 2019; Lynn et al., 2019), could be strategies to limit this risk.

## Material and Methods

### T cell sorting

CD4^+^ T cells were isolated from healthy adults using the RosetteSep human CD4^+^ T cell enrichment cocktail (STEMCELL Technologies, 15062) followed by a density gradient isolation (STEMCELL Technologies, 07861). Naïve Tregs were sorted from a CD25 enriched fraction ((Miltenyi, 130-092-983) CD4^+^CD127^lo^CD25^hi^ CD45RA^+^CD62L^hi^ cells and naïve CD4^+^ Tconvs were sorted from the CD25 negative fraction as live CD4^+^ CD45RA^+^CD62L^hi^ cells using a MoFlo® Astrios (Beckman Coulter). Cells were resuspended at 1×10^6^ cells/ml in XH media (X-Vivo 15 (Lonza) with 5% human serum, 1% penicillin/streptavidin, 1% glutamine and phenol red) containing 300U IL2/ml (Proleukin).

### Repetitive stimulation

sorted naive Tregs and Tconvs were stimulated with artificial antigen presenting cells (MacDonald et al., 2019a) loaded with αCD3 mAb (OKT3, UBC AbLab; 100ng/mL) in XH media with 300U IL2/ml, or stimulated with CD3/28 Dynabeads (ThermoFisher Scientific, 11141D) 3 bead: 1 cell ratio. Every week, a portion of cells was used for assays and the remaining were re-stimulated using the same initial stimulus with the exception that the Dynabead ratio was reduced to a 1 bead:1 cell ratio.

### Viral vectors and transduction

MSGV retroviral vectors and retroviral supernatant encoding the CD19-28z and gd2-28z CARs were produced as previously described (Lynn et al., 2019). For the retroviral transduction, sorted cells were stimulated with a 3 bead:1 cell ratio of Dynabeads in XH media containing 300U IL2/ml (Proleukin) as above. On the following day, non-tissue culture treated 96-well plates were coated overnight at 4 °C with 1 ml of retronectin (Takara) at 25μg/ml in PBS. On day 2 and 3, plates containing retronectin were washed with PBS and blocked with 2% BSA for 15 min. 100μl of thawed retroviral supernatant was then added to the well and centrifuged for 2h at 32 °C at 3,200 rpm before the addition of the same volume of media containing the activated cells. Media was refreshed on day 5. At day 7, cells were collected and stained with a fixable viability dye (FVD, Thermo Fisher Scientific), CD4 (clone RPA-T4, BD) and an anti-14g2a idiotype antibody (clone 1A7, NIH) or Protein L (genscript) to sort cells expressing the GD2-or CD19-specific CARs, respectively. Live CD4^+^CAR^+^ cells or live CD4^+^ cells (for the untransduced controls) were sorted (MoFlo® Astrios) resuspended at 0.5×10^6^ cells/ml in XH media containing 300U IL2/ml for an addition 4-5 days. Throughout cell culture, cell number and viability were assessed using the Cellometer auto2000 and an AOPI Staining solution (Nexelcom Bioscience).

### Flow cytometry

Cells were stained with fixable viability dye (FVD, Thermo Fisher Scientific, 65-0865-14) and surface markers in PBS for 30 min at 4°C before fixation and permeabilization using eBioscience FOXP3/Transcription Factor Staining Buffer Set (Thermo Fisher Scientific, 00-5523-00) and staining for intracellular proteins. Samples were analyzed on a BD LSR II (BD Bioscience) or a Cytoflex (Beckman Coulter) and results analyzed using FlowJo Software version 10.5.3. We used a FVD dye (Thermo Fisher Scientific, 65-0865-14), and the following antibodies from BD Biosciences; CD4 (RPA-T4), GARP (7B11), Ki67 (B56), CTLA4 (BNI3), CD8 (HIT8a), CD3 (UCHT1), IL-2 (MQ1-17H12) and an APC-Streptavidin conjugated antibody (554067). We also used CD25 (4E3, Miltenyi), and the following antibodies from eBioscience; CD127 (eBioRDR5), CD45RA (HI100), CD62L (DREG-56), FOXP3 (236A/E7), LAG-3 (3DS223H), CD4 (RPA-T4) and TNF-*α* (MAb11), IL-17A (eBio64DEC17). CD8 (RPA-T8), TIM-3 (F38-2E2), PD-1 (EH12.2H7), CD69 (FN50), IFN-*γ* (4S.B3) and Helios (22F6) were from Biolegend and GITR (110416) and LAP (27232) from R&D.

Apoptosis was assessed using the apoptosis/necrosis detection kit (Abcam, ab176749) according to the manufacturer’s instructions. For intracellular cytokine staining, 0.1×10^6^ cells were activated with PMA (10 ng/ml) and ionomycin (500 ng/ml) in the presence of Brefeldin A (10 μg/ml) for 4h at 37°C. Surface and intracellular staining was done as described above.

To monitor human cell engraftment in mice, 50 μL of blood was collected weekly and at the endpoint. The spleen was also collected at endpoint, weighed and 3-6μg used for staining. Red blood cells were lysed with ammonium chloride and remaining cells were resuspended in PBS containing anti-mouse CD16/32 (Thermo Fisher Scientific, 14-0161-82) for 10 minutes and then stained for extracellular markers using fixable viability dye (FVD; Thermo Fisher Scientific, 65-0865-14), anti-mouse CD45 (Thermo Fisher Scientific, 25-0451-82), anti-human CD45 (BD Biosciences, HI30), CD4 (Biolegend, RPA-T4), CD8 (eBioscience, HIT8a) as previously described (Dawson et al., 2019). Ten thousand counting beads were added to every sample (Thermo Fisher Scientific, 01-1234-42).

### TSDR

DNA was isolated and bisulfite-converted using the EZ DNA Methylation-Direct Kit (Cedarlane labs, D5021). A PCR of BisDNA was then made using the folowing primers: AGAAATTTGTGGGGTGGGGTAT (PCR FWD), ATCTACATCTAAACCCTATTATCACAACC (PCR REV-Bio) and AGAAATTTGTGGGGTGGG (SEQ FWD), followed by pyrosequencing using PyroMark buffers and assay on a Biotage PyroMark Q96 MD pyrosequencer (Qiagen). Methylation of the TSDR were calculated with the Pyro Q-CpG software (Biotage). Average methylation of the CpG sites 49260827, 49260817, 49260814, 49260808, 49260804, 49260796, 49260787 according to their position on the X chromosome (reference genome GRCh38) were reported.

### Seahorse Extracellular Flux assay

Seahorse XF Cell mito stress test assays were done using a Seahorse XFe96 Extracellular Flux Analyzer (Agilent) according to the manufacturer’s instructions. The day before, cartridge was hydrated using sterile water. The plate was coated with Cell-Tak solution (BD) and both the plate and 25 ml of XF calibrant (Agilent) were placed overnight at 37°C in a non-CO2 incubator. On the day of the assay, Cell-Tak was removed, washed with PBS and cells were counted and resuspended in Seahorse XF RPMI media, pH 7.4 (Agilent), supplemented with 1% FBS, 1mM glutamax and 10mM glucose. 2×10^5^ live cells/well were plated in triplicate when possible. Oligomycin (1 μM), FCCP (1.5 μM) and Rotenone/Antimycin A (1 μM) (all Sigma) were added as indicated in port A, B and C.

### Suppression assay

Allogeneic PBMCs were labeled with ef450 and Tregs with ef670 (ThermoFischer Scientific, 65-0842 and 65-0840, respectively). 0.1×10^6^ PBMCs were then plated in a 96-well plate with CD3/28 Dynabeads at a 1:16 cell: bead ratio. Labeled Tregs were added at the indicated ratios. For some experiments, nivolumab (Bristol Myers Squibb) was added at 10μg/ml. After 4 days, cells were stained extracellularly and CD4^+^, CD8^+^ and Treg proliferation was assess by flow cytometry. Suppression was determined as 100-(Division index with Tregs/Division index without Tregs *100).

### Xenogeneic graft-versus-host disease

8-to 12-week-old male and female NSG mice (NOD.*Cg-Prkdc*^*scid*^*Il2rg*^*tm1Wjl*^/SzJ, The Jackson Laboratory, bred in house) received whole-body irradiation (150 cGy, RS-2000 Pro Biological System), and one day later i.v. injected with PBS or 6 × 10^6^ PBMCs in the absence or presence of 3 × 10^6^ allogeneic untransduced or TS-CAR Tregs. Graft versus host disease was scored on the basis of weight, fur texture, posture, activity level, and skin integrity, with 0 to 3 points per category as described (Dawson et al., 2019). Mice were euthanized during the course of the experiment if they had a total score of 6 or a score of 3 in any category. Flow cytometric analysis to measure human immune cell engraftment is described above.

### RNA sequencing

RNA was extracted from 0.25×10^6^ cells at day 12 of cell culture using the Monarch total RNA miniprep kit (New England Biolabs, T2010S) and quantified using the Qubit RNA HS Assay kit (Thermo Fisher Scientific, Q32855) according to the manufacturer’s instructions. RNA purity (RNA Integrity Number) was assessed using the RNA 6000 Pico assay (Agilent, 5067-1513) and samples prepared using the TruSeq stranded mRNA library kit (Illumina) on the Illumina Neoprep automated nanofluidic prep instrument. Illumina NextSeq 500 with Paired End 42bp × 42bp reads was used for sequencing and de-multiplexed read sequences and sequenced were then aligned to the Homo sapiens (PAR-masked)/hg19 reference using STAR aligner (version 2.5.0a) and the RNA-Seq Alignment App (version 1.1.0) on Illumina Basespace. De-multiplexed read sequences and RnaReadCounter (part of the internal analysis tool IsisRNA (version 2.6.25.18)) was used for counting the number of aligned reads, as described previously (Hoeppli et al., 2019). In R, raw count matrices were generated using HTSeq (v0.11.2), then scale factors were calculated to take into account differences in library sizes using edgeR (v3.24.3) and normalization was performed using limma (v3.38.3) as in (Law et al., 2016). Log (CPM) and visualization was performed using: ggplot2 (3.2.1), RColorBrewer (v1.1.2), tibble (2.1.3), pheatmap (v1.0.12), stats (v3.5.1), and gplots (v3.0.1.2). The Gene Set Enrichment Analysis was performed with fgsea (v1.8.0) and Hallmark gene sets from MSigDB (v6.2). The code used for data analysis is available on GitHub: https://github.com/fransilvion/RNA_seq_Lamarche. RNAsequencing data are deposited on NCBI GEO, GSE153384.

### Statistical analysis

All statistics were done using Prism version 8.4.2 and are described in the figure legends. For the experiments where multiple assays were repeated over time (Figure 1), one-way ANOVA or mixed-effects analysis with Dunnett’s multiple comparisons test were used, comparing each data to their matched results at day 7. To compare the different Tregs, one-way Anova with Turkey’s comparisons test were done. **** p ≤ 0.0001, *** p ≤ 0.001, ** p ≤ 0.01, * p ≤ 0.05.

### Study Approval

Healthy volunteers gave written informed consent to use their blood according to protocols approved by the University of British Columbia Clinical Research Ethics Board (UBC-CREB) and Canadian Blood Services. Animal protocols were approved by the UBC Animal Care Committee.

## Author contributions

CL conceived, designed and performed experiments, analyzed data and wrote the manuscript; GEN carried out bioinformatic analysis and critically reviewed the manuscript; CNQ conducted experiments and analyzed data; EWW and CLM provided essential reagents, intellectual input and critically reviewed the manuscript; MKL conceived and designed experiments, provided overall direction and interpretation, obtained funding and wrote the manuscript.

## Acknowledgments

This work was supported by a grant from the Canadian Institutes of Health Research (CIHR) FDN-154304. CL is supported by a CIHR fellowship and a KRESCENT fellowship award. MKL receives a salary awards from the BC Children’s Hospital Research Institute. We thank Dr. Ramon Klein-Geltnick for advice and critical reading of the manuscript and Dr. Lixin Xu for expert flow cytometry support. MKL received research funding from TxCell, Bristol-Myers Squibb, Pfizer, Takeda and CRISPR Therapeutics for work unrelated to this report. CLM holds several patent applications in the area of CAR based immunotherapy, and is a founder of, holds equity in, and receives consulting fees from Lyell Immunopharma which develops CAR based therapies. EWW holds several patent applications in the area of CAR based immunotherapy and receives consulting fees from Lyell Immunopharma. CLM is a member of the Parker Institute for Cancer Immunotherapy, which supports the Stanford University Cancer Immunotherapy Program. The authors have no additional financial interests.

## Abbreviations

CAR: Chimeric antigen receptor
ECAR: extracellular acidification rate
OCAR: oxygen consumption rate
Tconv: conventional T cell
Treg: Regulatory T cell
TS: tonic signaling
TSDR: Regulatory T-cell-Specific Demethylated Region

