## Supplemental Figures for "Repeated stimulation or tonic-signaling chimeric antigen receptors drive regulatory T cell exhaustion"

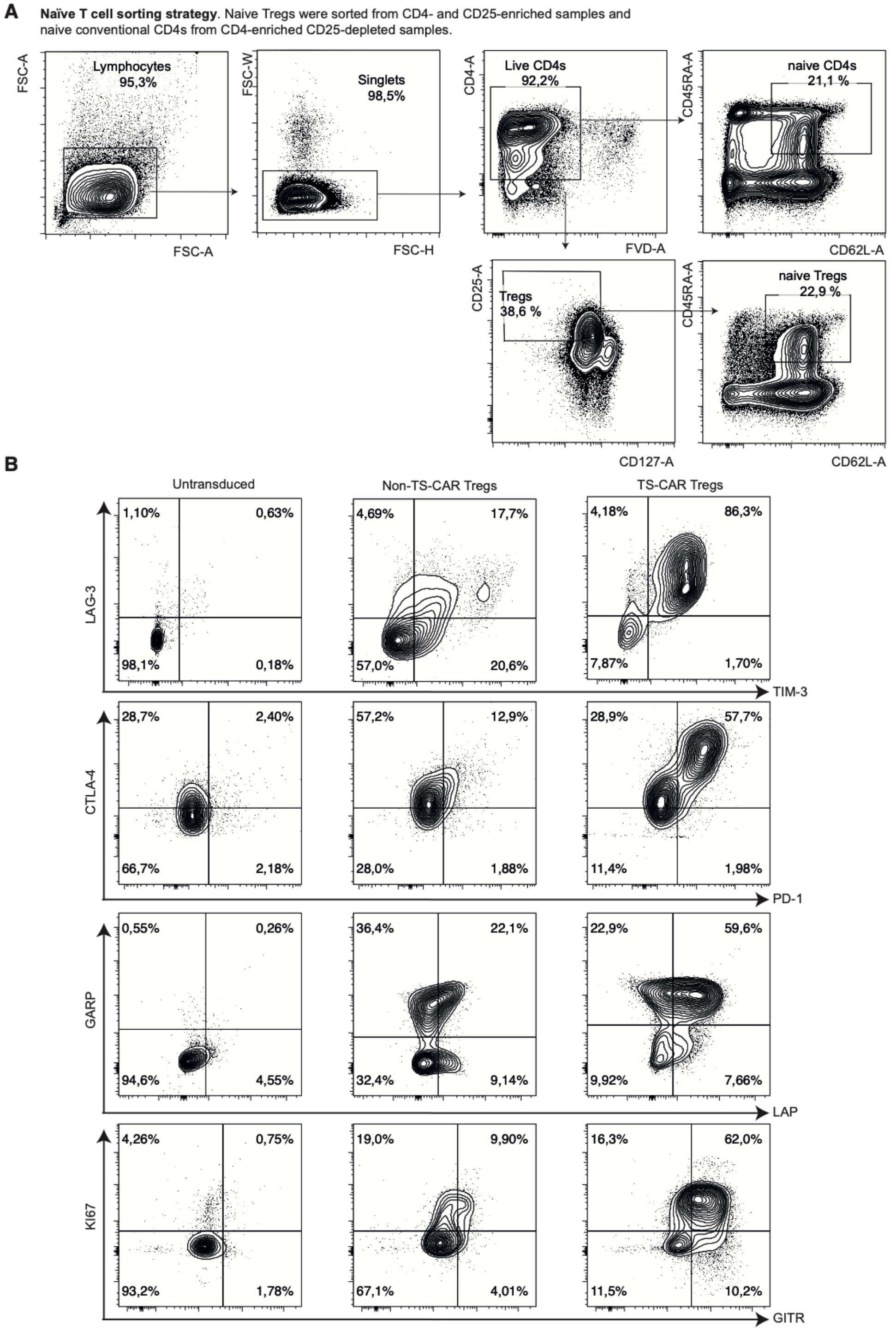


**Supp Figure 1.** ***Cell sorting strategy and representative flow cytometry data.*** **(A)**. CD4$^{+}$ T cells were enriched from PBMCs using a RosetteSep human CD4 enrichment cocktail and were then separated into CD25-enriched and -depleted fractions using magnetic separation. Naïve Tregs were sorted as live CD4$^{+}$CD127$^{lo}$CD25$^{hi}$ CD45RA$^{+}$CD62L$^{hi}$ cells from the CD25-enriched fraction and naïve CD4$^{+}$ Tconvs as live CD4$^{+}$ CD45RA$^{+}$CD62L$^{hi}$ cells from the CD25-depleted fraction. **(B)** Representative flow cytometry data for surface receptors (LAG-3, TIM-3, CTLA-4, PD-1, LAP, GARP and GITR) and KI67, comparing untransduced Tregs to Tregs transduced with a non-tonic signaling (non-TS) or a tonic-signaling (TS)-CAR.

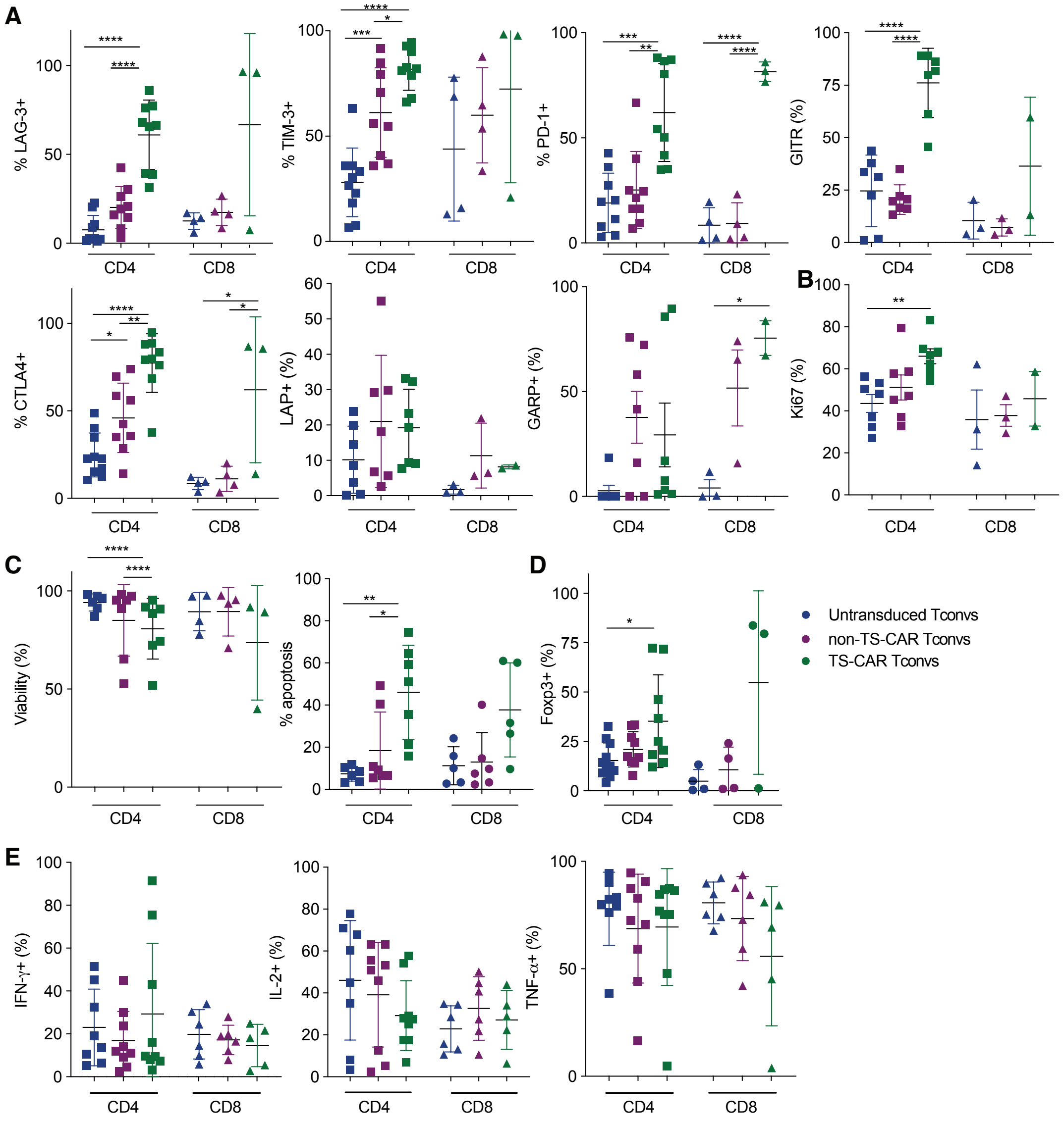


**Supp Figure 2.** ***Expression of a tonic signaling CAR in CD8⁺ and CD4⁺ Tconvs induces phenotypic changes consistent with exhaustion.*** Naïve $CD4^+$ and $CD8^+$ T cells were left transduced (UT) or transduced with retrovirus encoding a non-tonic signaling (non-TS) or a tonic signaling (TS-) CAR. After 11-12 days of culture cells were analyzed for expression of **(A)** LAG-3, TIM-3, PD-1, GITR, CTLA4, LAP and GARP, **(B)** Ki67. **(C)** Viability was assessed using an automated cell counter, apoptosis by flow cytometry and **(D)** FOXP3 by intracellular staining. **(E)** Intracellular cytokine production (IL-2, IFN-γ and TNF-α) were assessed by flow cytometry 4 hours after PMA/Ionomycin stimulation. Each dot is a unique donor. Mean ± SD. One-way Anova with Turkey's comparisons test were done for each of the comparisons. **** $p \leq 0.0001$, *** $p \leq 0.001$, ** $p \leq 0.01$, * $p \leq 0.05$.

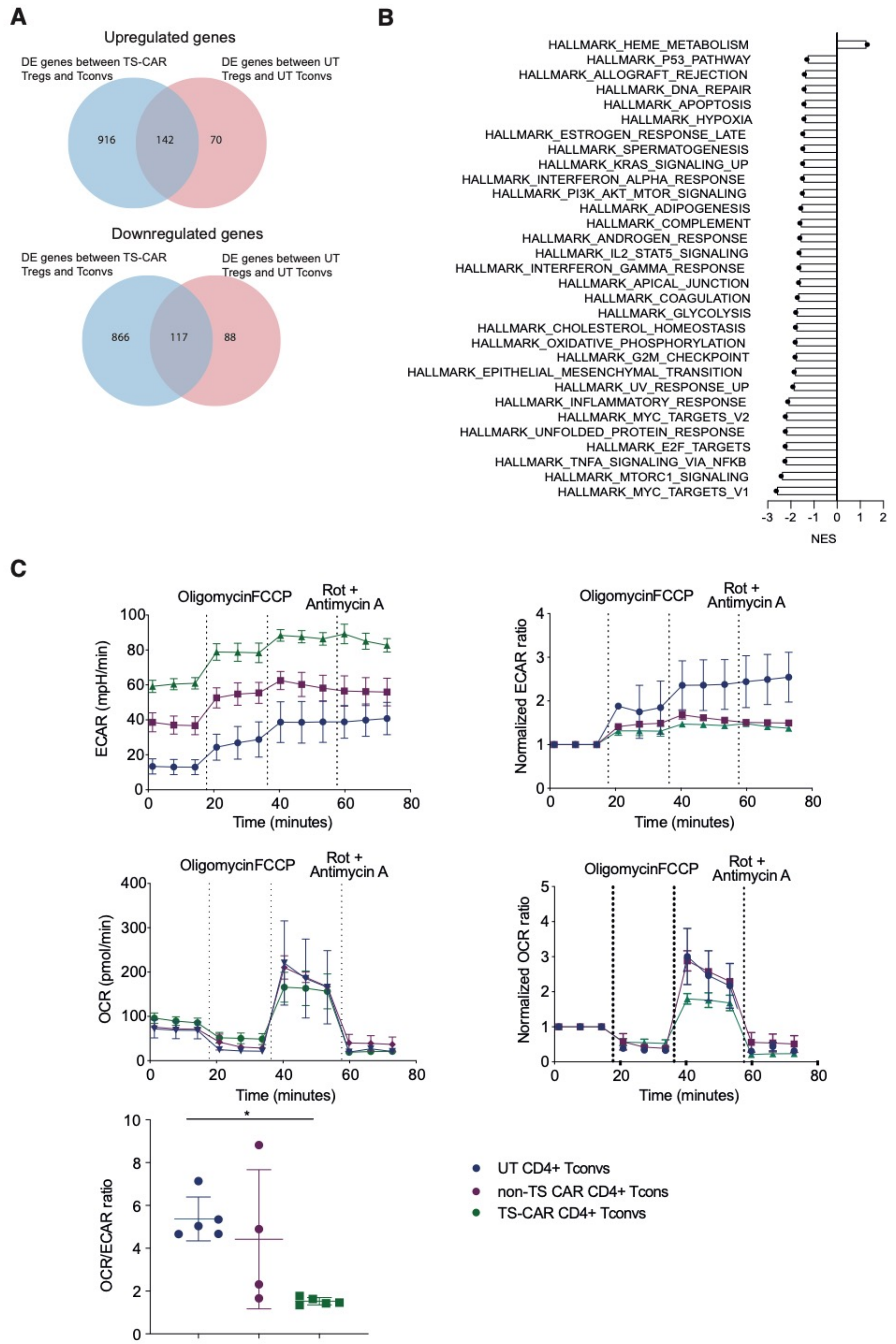


**Supp Figure 3.** ***Expression of a tonic signaling CAR induces changes in the transcriptome and metabolism.*** **(A&B)** After 12 days of culture, RNA was extracted from Tregs or CD4$^+$ Tconvs that were untransduced (UT) or transduced to express a non-tonic signaling (non-TS) or a tonic signaling (TS) CAR, and subject to RNA sequencing. (**A**) Venn diagram showing the number of differentially expressed (DE) genes between TS-CAR Tregs and Tconvs (blue) and between UT Tregs and Tconvs (red), either upregulated (top) or downregulated (bottom). Genes included in **Figure 3C** were DE between TS-CAR and Tconvs but not between UT Tregs and Tconvs (916 upregulated and 866 downregulated). (**B**) Gene set enrichment analysis within the indicated types of $CD4^+$ Tconvs. Normalized enrichment score (NES) of hallmark pathways analysis overrepresented in TS-CAR Tregs compared to TS-CAR $CD4^+$ Tconvs. n=3 per group from 2 independent experiments. **(C)** After 11-12 days of culture, the indicated types of $CD4^+$ Tconvs were plated in Cell-Tak coated wells and subject to the Agilent Seahorse XF Cell Mito Stress Test. Extracellular acidification rate (ECAR) (top left) and normalized ECAR on baseline (top right) over time, Oxygen consumption rate (OCR) (middle left) and normalized to baseline (middle right). Oligomycin, FCCP and a mix of Rotenone and antimycin were added as indicated. n=2-3 donor from 2 independent experiments, using the average of technical replicates. (bottom) OCR/ECAR ratio at baseline n=4-5 from 3 independent experiments. One-way Anova with Turkey's comparisons test, p=0.0162. Mean ± SEM.
